# RNA modification mapping with JACUSA2

**DOI:** 10.1101/2021.07.02.450888

**Authors:** Michael Piechotta, Qi Wang, Janine Altmüller, Christoph Dieterich

**Affiliations:** Klaus Tschira Institute for Integrative Computational Cardiology, University Hospital Heidelberg, 69120 Heidelberg, Germany; Cologne Center for Genomics (CCG), University of Cologne, Weyertal 115b, 50931 Köln, Germany; Department of Internal Medicine III (Cardiology, Angiology, and Pneumology), University Hospital Heidelberg, 69120 Heidelberg, Germany; German Centre for Cardiovascular Research (DZHK)-Partner Site Heidelberg/Mannheim, 69120 Heidelberg, Germany

**Keywords:** m6A, Pseudouridine, Reverse transcription signature, Nanopore, Multi-threading

## Abstract

A whole series of high-throughput antibody-free methods for RNA modification detection from sequencing data emerged lately. We present JACUSA2 as a versatile software solution and comprehensive analysis framework for RNA modification detection assays that are based on either the Illumina or Nanopore platform. Importantly, JACUSA2 can integrate information from multiple experiments (e.g. replicates and different conditions) and different library types (e.g. first- or secondstrand libraries). We demonstrate its utility by example, showing three analysis workflows for m6A detection on published data sets: 1) MazF m6a-sensitive RNA digestion (FTO+ vs FTO-), 2) DART-seq (YTHwt vs YTHmut) and 3) Nanopore profiling (*METTL3* +/+ vs -/-). All assays have been conducted in HEK293 cells and complement one another.

## Background

Lately, we have seen a whole wave of antibody-free methods to map RNA modifications transcriptome-wide based on high-throughput sequencing data (1;2). What is common to all of these approaches is that, independent of the actual protocol, RNA modification sites manifest as specific read signatures in sequencing data. Others and we have previously worked on mapping modifications, which originate from RNA editing (3). For example, adenosine deamination is a process that typically converts A *→* I and is facilitated by the family of ADAR proteins (4). Previously, we had contributed JACUSA (5) to this exciting field of RNA modification research.

In this manuscript, we present the successor JACUSA2 (https://github.com/dieterich-lab/JACUSA2), which features better running time performance and captures more complex read signatures involving base substitutions, insertions, deletions, and read truncations (see Figure 1A-C). Novel antibody-free approaches have been established for the mapping of N6-methyladenosine (m6A), 5-methylcytosine (m5C), pseudouridine (Ψ) and N1-methyladenosine (m1A) and beyond by sequencing (reviewed in (6)). In this manuscript, we focus on messenger RNA (mRNA) modifications in general and N6-methyladenosine (m6A) in particular. However, our software approach generalizes to other RNA species (see Supplementary Text for an rRNA example) and all of the aforementioned modifications as the presence of a modified residue typically manifests as a site-specific increase in 5’ or 3’ read termination, base misincorporation, and insertion or deletion rate. An additional feature is the possibility to stratify reads by a particular base substitution, which could occur anywhere on the read. JACUSA2 is able to process data from any read technology as long as it can be provided in the binary sequence alignment map format (BAM), a universal standard format (7). In the following, we have applied JACUSA2 to sequencing data from the Illumina and ONT Nanopore platform across 3 use cases. Downstream analyses of JACUSA2 output are supported by an additional R package JACUSA2helper (https://github.com/dieterich-lab/JACUSA2helper)

**Figure 1:**
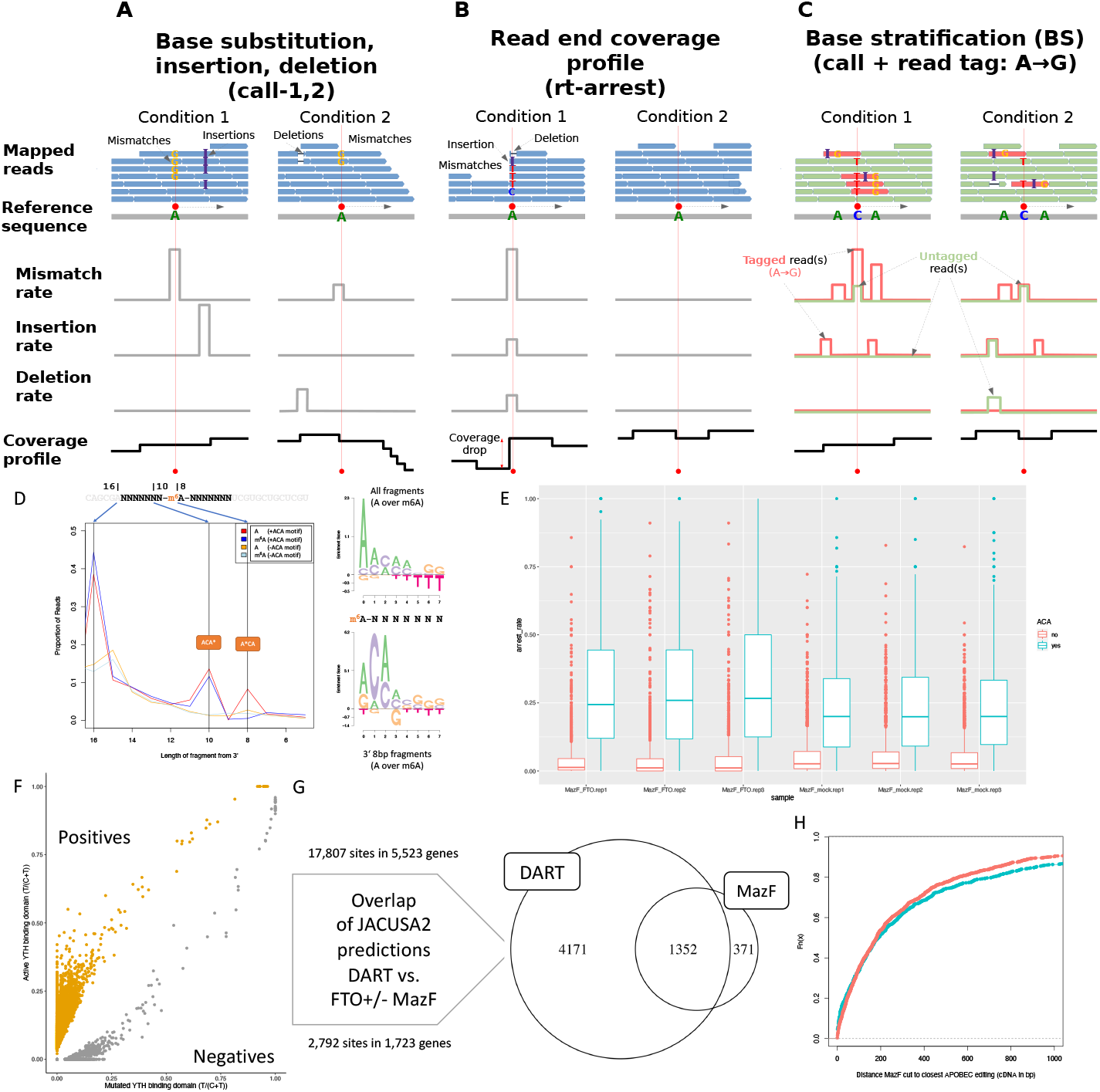
Summary of JACUSA2 analysis options and illustration of utilized sequencing data properties. All analysis options can be extended with INDEL calling where insertion and/or deletions rates between 2 conditions are modelled to identify sites with statistically significant divergent INDEL rates. The matching JACUSA2 command line option is given in parenthesis, e.g.: (call) for variant calling. A: Variant calling: Modelling and comparison of mismatch rates between 1 or 2 conditions. B: Arrest site detection: Coverage profiles of two conditions are modelled to discover statistically significant coverage drops that are termed arrest site. C: Base stratification: Variant calling can be extended with base stratification where reads are grouped or tagged by a specific base substitution. This enables to compare tagged and untagged reads between conditions. D: Synthetic RNA Oligo digestion and sequencing. Position 8 hosts the central A/m6A. Position 1-7 and 9-16 are random. The MazF recognition site is derived from this assay. E: Arrest rate analysis across replicates in published data from HEK-293 cells. We define a set of 2,792 well covered sites with robust and significant read arrest changes. F: Comparison to DART-seq analysis, which uses an m6A reader domain fused to APOBEC1 (C *→* U). We overlap 17,087 JA-CUSA2 positive predictions from DART-seq and FTO-MazF on gene and individual site level. G: Overlap of predicted gene sets from MazF (1,723 8 modified gene loci) and DART (5,523 modified gene loci). H: cDNA distance of closest DART signal (C *→* U) to each differential MazF cleavage side.

## Results and Discussion

We present three representative use cases in RNA modification detection on mRNA. The first use case focuses on a methylation-sensitive RNA restriction enzyme (MazF) assay to identify m6A modified residues in mRNA of HEK293 cells (8; 9). The second use case complements the first by demonstrating a genetic approach to map m6A sites in HEK293 cells using DART-seq (10). While all of the aformentioned use cases utilized Illumina sequencing data, the last use case analyzes Nanopore sequening data from a HEK293 wildtype and knockout cells (*METTL3* -/-) (11).

### Use Case 1: m6A mapping by MazF treatment and sequencing

Lately, we have seen a whole wave of antibody-free methods to map m6A RNA modifications transcriptome-wide based on high-throughput sequencing data. A need for antibody-free complementary approaches has been clearly identified by a rigorous assessment of published MeRIP data sets. McIntyre et al. show that m6A peak overlap varies from *≈* 30 to 60% between studies, even in the same cell type (12). An antibody-independent solution to map m6A sites is provided by an alternative biochemical approach, namely m6A methylation-sensitive enzymatic RNA digestions (8; 9). We have tested the substrate specificity of the MazF enzyme on custom RNA oligos with a centrally modified A position. Figure 1D gives an overview on the taken approach to assess MazF substrate specificity. Briefly, the central A/m6A position is flanked by 7 random nucleotides on each site and a constant prefix and suffix. Total sequence length per oligo is 34bp. These RNA oligos were subjected to enzymatic digestion. As previously reported, MazF cuts 5’ to an ACA motif. Cutting efficiency is reduced by the presence of an m6A modification (A*CA motif, asterisk denotes modification). This enzymatic reaction does not differ for cutting 5’ of ACA or ACA*. We have reanalyzed the 6 Illumina libraries from Zhang et al. (8). Briefly, 3 replicates were treated with FTO, a demethylase that removes m6A from mRNA (13), and 3 replicates were mock-treated. Figure 1E shows the observed arrest rates in the data stratified by the presence or absence of an ACA motif. Arrest rates are significantly elevated in FTO-treated samples as compared to mock-treated samples. This characteristic read signature is used to predict m6A modified sites in mRNA in a head-to-head comparison using JACUSA2 (FTO vs WT).

### Use Case 2: DART-seq

A complementary genetic approach is an extension of the TRIBE technique called DART-seq (10). Meyer applied DART-seq on HEK293 cells where the APOBEC domain was fused to the YTH domain from human YTHDF2 (WT and mutated). In essence, new C*→*U editing events that are significantly enriched in the YTHDF2-WT, but not in the binding domain mutant are bona fide candidates for m6A RNA modification. Figure 1F shows the estimated cDNA base substitution rates T/(C+T) for the active and mutated YTH binding domain. Elevated C *→* U deamination on RNA level for the active domain over the mutated domain are used to predict m6A modifications on mRNA. Generally, DART-seq has the advantage that it can be extended to 3rd generation long read sequences and requires only very little RNA input amounts (*in vivo* experiments) yet relies on genetic modification of the target system / organism. Figure 1G summarizes the analysis results and identifies 5,894 gene loci, which produce potentially m6A-modified mRNAs (union of use case 1 + 2). The overlap between both approaches is 1,352 gene loci. Figure 1H shows the genomic distance of a called C *→* U site to the closest differential MazF digestion site.

### Use Case 3: Nanopore direct RNA sequencing

Direct RNA sequencing without involving any cDNA library preparation or PCR-based amplification steps is a unique feature of the ONT sequencing platform. Briefly, native poly-adenylated RNA is threaded through a nanopore in 3’ *→* 5’ orientation (Figure 2A). Some ground-breaking work on m6A detection by direct RNA sequencing has been performed by Liu et al. (14) in yeast and Jenjaroenpun et al. (15) in mouse embryonic stem cells. We have identified a published direct RNA sequencing data set from HEK293 cells (11), which facilitates a direct comparison to the aforementioned Illumina-based approaches. Pratanwanich et al. use a genetic ablation of *METTL3* and contrast read error profiles and signal traces to wildtype (reference) samples.

**Figure 2:**
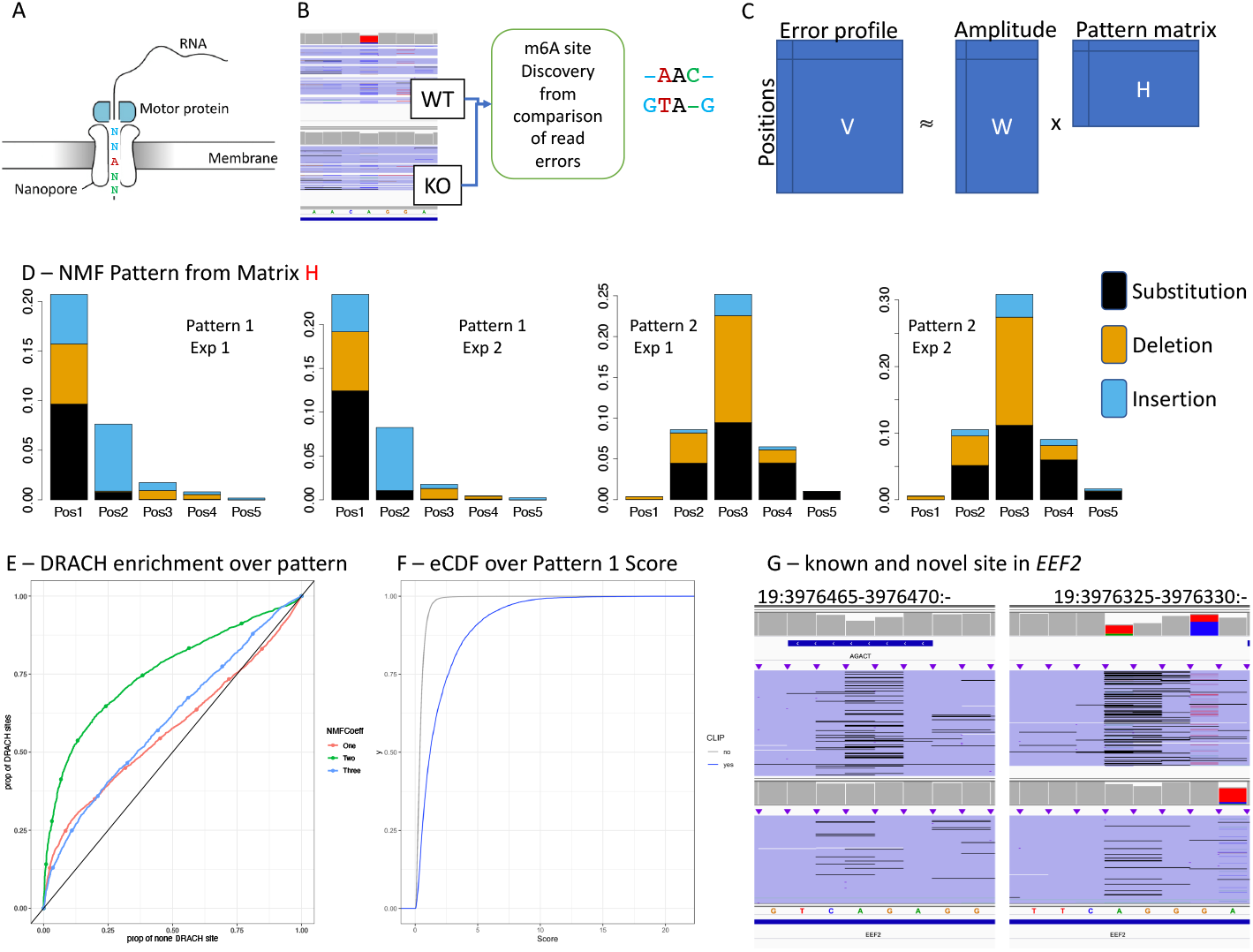
Nanopore sequencing of HEK293 cell lines: Mettl3 KO and WT. A: Direct RNA sequencing on Nanopore. B: Signature detection with JACUSA2 comparing WT over KO cells. Three principal events are detected: base substitutions, insertions and deletions. All 5mers with a central A (NNANN) are considered. C: Non-negative matrix factorization to identify characteristic patterns (matrix H) that are indiciative of the m6A modification. D: Two main patterns (out of 5). The first pattern coincides with the occurrence of the DRACH motif. E: Pattern 1 is enriched for the DRACH motif. 5mer occurences are ranked by their pattern strength. The proportion of DRACH over non-DRACH motifs is plotted. F: Pattern 1 score distribution stratified by miCLIP validated sites and other sites. G: identification of a known and novel site in the *EEF2* gene locus.

We applied JACUSA2 to identify read error profile differences between the respective paired samples (see Figure 2B). We are considering all positions with sufficient read coverage (*>*4) across all replicates and focus on positions with a specific 5mer context, which is basically a central A nucleotide flanked by 2 adjacent random nucleotides (NNANN). Our initial unsupervised pattern learning approach starts from 8,983 previously reported m6A sites (16).

Only non-overlapping 5mers outside of homo-polymer regions (JACUSA filter: Y) are subsequently used. The respective feature set consists of deletion, insertion, and substitution scores for every considered 5mer. This amounts to a feature matrix of 3 event scores *×* 5 motif positions *×* 2 experiments = 30 features. We subjected this feature matrix to a non-negative matrix factorization (NMF) analysis (Figure 2C). Through robustness and concordance assessment, we could identify an optimal factorization rank of 5. The pattern matrix (H) highlights the importance of pos 3 (and 4) for the most relevant pattern (i.e. pattern 1). The loading of this factor correlates with the presence of the well known DRACH motif (Figure 2D). The cumulative density function of pattern 1 scores can then be used to predict novel m6A sites from *>* 2 million covered candidate sites (NNANN, Figure 2E). This is exemplified by 2 high-scoring m6A sites in the *EEF2* gene locus (one known, one novel, Figure 2F).

## Conclusion

We present JACUSA2 as a versatile software solution and comprehensive analysis framework for RNA modification detection assays that are based on either the Illumina or Nanopore platform. We have explored three state-of-the art use cases for m6A detection in mRNA. All of which use antibody-free methods and encompass complex read signatures. JACUSA2 supersedes our initial software in terms of features and speed (see SFig 1). A summary table of m6A site identifications is provided as supplementary table (STab 1). In the Supplementary Material, we highlight an additional use case (Ψ detection on human rRNA transcripts) outside of mRNA modification analysis (see SFig 2 and STab 2). JACUSA2 is complemented with an R package (JACUSA2helper), which eases downstream processing of JACUSA2 output (https://github.com/dieterich-lab/JACUSA2helper). We provide all relevant information (data and code) to reproduce our results from the presented workflows as vignette in the JACUSA2helper repository. We will continue to develop JACUSA2 to meet new requirements in terms of prediction performance for all aforementioned sequencing platforms.

## Availability of data and materials

A comprehensive list of all sequencing data sets can be found in Supplementary Materials. Our software can be either found at: the JACUSA2 repository (JAR files + tutorials, https://github.com/dieterich-lab/JACUSA2) or the JACUSA2helper repository (R package + vignettes, https://github.com/dieterich-lab/JACUSA2helper)

## Author’s contributions

MP designed and implemented JACUSA2 and JACUSA2helper. QW tested the software and implemented use cases. CD acquired funding, supervised the project and analyzed data. CD and MP wrote the manuscript. JA helped with MazF assay development and sequencing

## Acknowledgements

We would like to thank all members of the Dieterich Lab for valuable discussions. We are grateful to Christian Becker and Marek Franitza at the Cologne Center for Genomics (CCG) for their invaluable library preparation and sequencing support. We thank Volker Boehm for discussing an early attempt of the MazF digest with us.

## Figures Additional Files

**Additional file 1 — Supplementary Text (PDF)**

**Additional file 2 — Supplementary Table 1 (XLSX)**

**Additional file 3 — Supplementary Table 2 (XLSX)**

