## Supplementary Document for "RNA modification mapping with JACUSA2"

July 2, 2021

#### Methods

The variant calling framework from JACUSA1 has been redesigned to enable efficient read processing beyond the identification of single nucleotide variants (SNVs). Native support for single and paired end reads for common stranded and unstranded library types has been added and the core BAM processing component has been upgraded to htslib (`https://github.com/samtools/htsjdk`). We designed a benchmark to compare the running time performance between different implementations based in the variant calling setup (check Figure S1 for detailed results).

A new analysis mode (*rt-arrest*) has been added to JACUSA2 that allows to study read arrest events in paired samples. This works even for combining different library types (see Figure 1B in main text). Furthermore, JACUSA2 allows to identify differential insertion and/or deletion sites. Finally, variant stratification has been added as an optional feature to study the co-occurence of a specific variant and other features such as variants, arrest sites, or INDELs (see Figure 1C in main text). JACUSA2helper (`https://github.com/dieterich-lab/JACUSA2helper`) complements JACUSA2 to study RNA modifications. It is a R package that simplifies downstream analysis and visualization of JACUAS2 output.

##### General JACUSA2 analysis workflow

In general, JACUSA2 features a two tier analysis:

**JACUSA2 (Java)** fast and accurate variant or arrest event detection and

**JACUSA2helper (R)** detailed followup analysis and visualization of results.

Given mapped reads as input (BAM files) and corresponding library type information, JACUSA2 will assign aligned reads to separate sequence coordinate windows. Each window is handled by separate threads (i.e. independent set of reads), which enables fast parallel computation on modern machines. Reads are filtered according to user provided parameters, and processed according to the chosen method.

JACUSA2 supports the following main analysis methods:

- simple pileup (*pileup*) of mapped reads from arbitrary number of conditions
- variant calling (*call-{1,2}*) with one or two conditions
- read arrest detection (*rt-arrest*) with two condition

Each method features a statistical test that provides a test-statistic to filter significant candidates. Additionally, each analysis method can be extended to identify differential INDELs and/or to stratify observations based on a specific co-occuring variant.

Optional artifact/feature filtering can be carried out to remove artifacts that cannot be reliably captured by the statistical model (e.g., homopolymers). Finally, results from each threaded window are collected and combined into the final output.

#### Variant calling

The existing variant calling framework in JACUSA1 has been reimplemented for JACUSA2 and variant calling for only one condition has been added. In this single condition variant calling mode, an *in silco* condition is created from the sequencing data by transforming non-reference bases to conform with the reference sequence. A reference FASTA can be provided by the user or the MD-field from the SAM File Format is used to reconstruct the reference sequence.

#### General modelling framework

In order to model counts beyond base calls observed in variant calling, we revised the original implementation of the Dirichlet-Multinomial distribution $\boldsymbol{x} \sim DirMult(\alpha_1, \ldots, \alpha_K)$ from JACUSA1. The goal was to model category count vectors $\boldsymbol{x} = (x_1, \ldots, x_K)$ for arbitrary $K$.

In variant variant calling, the category count vector $\boldsymbol{x}$ consist of 4 elements and is defined as the observed base call counts $\boldsymbol{x_{BC}} = (x_A, x_C, x_G, x_T)$ at an arbitrary site within the genome. The Dirichlet Multinomial $DirMult(\boldsymbol{\alpha})$ is used to model base call counts $\boldsymbol{x_{BC}}$ and the Newton iteration method presented by Minka (https://tminka.github.io/papers/minka-newton.pdf) is used to determine parameters $\boldsymbol{\alpha}$.

We implemented a general framework based on the Beta-Binomial distribution to model counts of binary features derived from reads. The Beta-Binomial distribution $BetaBin(\alpha, \beta)$ can be defined as a special case of the more general $DirMult(\alpha_1, \ldots, \alpha_K)$ when $K = 2$. Given some arbitrary binary feature $y$ the count vector is defined as $\boldsymbol{y} = (y_1, y_2) : y = y_1 + y_2$. To account for zero counts and where appropriate, we add a pseudocount of one.

For each observed and pooled count vectors $BetaBin(\alpha, \beta)$ distributions are fitted with the aforementioned method and compared with a likelihood ratio test. The null hypothesis is that the counts of each condition $(c \in I, II)$ have the same underlying distribution:

$$LL = -2 \cdot \log \frac{BetaBin(\alpha^I, \beta^I) \cdot BetaBin(\alpha^{II}, \beta^{II})}{BetaBin(\alpha^{I+II}, \beta^{I+II})}$$

We approximate $LL$ with a $\chi^2$ distribution and provide the pvalue.

##### Modelling read arrest events

An arrest event is identified by comparing stop and read through events counts between two conditions $(c \in \{I, II\})$. Given the general modelling framework, the category count vector is defined as the number of observed **A**rrest events and read **T**hrough events $y_{AT} = (y_{arrest}, y_{through})$.

The arrest rate $r$ is defined as the fraction of arrest reads and total reads:

$$r_{arrest} = \frac{y_{arrest}}{y_{arrest} + y_{through}}$$

We model $\boldsymbol{y_{AT}}$ with the Beta-Binomial distribution and use the aforementioned general framework to perform a statistical test. We report the test-statistic and the approximated pvalue for each candidate arrest site.

##### Modelling INDELs

The underlying statistical framework of INDEL calling is comparable to the previously presented modelling of read arrest events. The general modelling framework is used to model observed insertion and/or deletion counts. In insertion calling, the category vector $\boldsymbol{y_{I\overline{I}}}$ is defined as the number of observed reads with $(y_{ins})$ and without $(y_{\overline{ins}})$ an insertion: $\boldsymbol{y_{I\overline{I}}} = (y_{ins}, y_{\overline{ins}}) : y_{\overline{ins}} = y_{tot} - y_{ins}$. The total number of reads $(y_{tot})$ includes all reads that span the respective insertion, including intron regions and low quality base calls. For each insertion site the category vector $\boldsymbol{y_{I\overline{I}}}$ is modelled with the Beta-Binomial distribution. The test-statistic and the approximated pvalue for each insertion site are reported.

Deletion calling works analogously to insertion calling with the category vector defined as: $\boldsymbol{y}_{D\overline{D}} = (y_{del}, y_{\overline{del}})$. Similar modelling and reporting applies here.

#### Variant stratification

In JACUSA2, a user customizable base substitution can be set to split reads into two categories. Reads that contain the base substitution of interest are termed tagged and the remaining set of reads is called untagged. This variant stratification enables to study the co-occurrence of a variant and some other feature such as arrest events, other variants, or INDELs (see Figure 1C - main text). Variant stratification requires stranded library types and supports single and paired end reads. In paired end reads, a read is termed tagged, if either of the two reads in the pair contains the variant.

### Comparison with JACUSA1.*

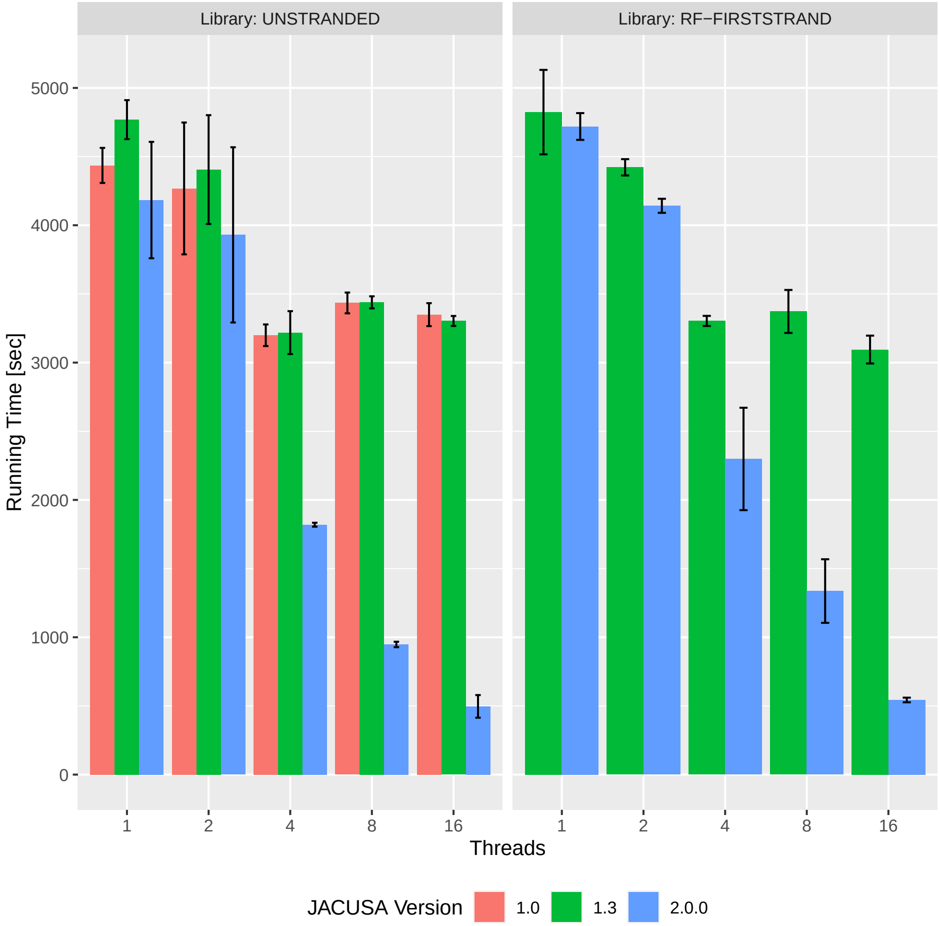


Figure 1: Average running time comparison of different JACUSA versions with unstranded vs. stranded library type (RF-FIRSTSTRAND) and increasing number of used CPU cores (threads). Tested on reduced BAM files (chr1 and chr2) from ADAR knockdown and untreated conditions from [1]. Each run has been repeated 3× (whiskers indicate fastest and slowest running time).

In brief, the benchmark aims to explore the influence of sequence library type (stranded vs. unstranded) and the number of utilized parallel threads on the running time. We used BAM files restricted to chr1 and chr2 from ADAR knockdown and untreated conditions from [1] as test data to identify varaints between both conditions. On average, the improved JACUSA2 implementation outperforms its predecessors in both unstranded and the more demanding stranded library type setting. More importantly, JACUSA2 scales much better showing a steady decrease of running time when the number of additional CPU cores is increased. In contrast, increasing the number of threads beyond 4 does not show any running time gains for JACUSA1.x. Utilizing 16 cores JACUSA2 completes the unstranded task on average after $\approx$ 500 seconds compared to $\approx$ 3.300 seconds for the previous versions. Overall, a significant running time enhancement can be observed when comparing against previous JACUSA versions (see Figure S1).

#### JACUSA 1.0

JACUSA 1.0 was downloaded and built from the "paper" branch: `https://github.com/dieterich-lab/JACUSA/tree/paper`. JACUSA 1.0 does not support stranded library types, therefore we have only evaluated performance in the unstranded scenario.

The following line was executed to run JACUSA 1.0:

```
$ java -jar JACUSA_v1.0.jar call-2 $OPTS -r $OUTPUT $INPUT
```

#### JACUSA 1.3

JACUSA 1.3 was downloaded from: `https://github.com/dieterich-lab/JACUSA/releases/download/1.3.0/JACUSA_v1.3.0.jar`.

The following line was executed to run JACUSA 1.3:

```
$java -jar JACUSA_v1.3.0.jar call-2 -P $LIB,$LIB $OPTS -r $OUTPUT $INPUT
```

#### JACUSA 2

JACUSA 2 was downloaded from: `https://github.com/dieterich-lab/JACUSA2/releases/tag/v2.0.0`.

```
$ java -jar JACUSA_v2.0.0.jar call-2 $OPTS -r $OUTPUT $INPUT"
```

#### Additional Use Case: Pseudouridylation site mapping in human rRNAs

Pseudouridylation ($\Psi$) is an abundant and widespread type of RNA epigenetic modification in living organisms. Specifically, rRNAs and tRNAs are well characterized targets of Pseudouridylation [2, 3]. In rRNA, pseudouridylation is carried out by ribonucleoprotein (RNP) complexes called H/ACA box RNPs, each consisting of one H/ACA snoRNA and four core proteins, namely GAR1, NHP2, NOP10 and dyskerin (DKC1) [4]. The recent identification of variable pseudouridylation sites has overturned the notion that all rRNA modifications are constitutively present on ribosomes, highlighting nucleotide modifications as an important source of ribosomal heterogeneity [5]. There are several ways to make $\Psi$ sites accessible to sequencing-based profiling. $\Psi$ sites have been mapped transcriptome-wide by chemically modifying pseudouridines with carbodiimide and detecting the resulting reverse transcription stops in high-throughput sequencing [6, 7, 8]. Specifically, $\Psi$ sites on the human 80S rRNA are well annotated [2] and CMC-based rRNA profiling data from different cDNA libraries are available [6]. For our particular use case, we compiled a list of 104 $\Psi^+$ sites and 1,061 $\Psi^-$ sites (i.e. other uridine residues), which are covered in all respective sequencing libraries (Figure 2A, Supplementary Table 2). We consider read truncation, misincorporation, insertions and deletions from 3 experimental conditions as a 24-dimensional feature set to identify $\Psi^+$ sites (3 experiments $\times$ 4 feature $\times$ 2 positions (0,+1)).

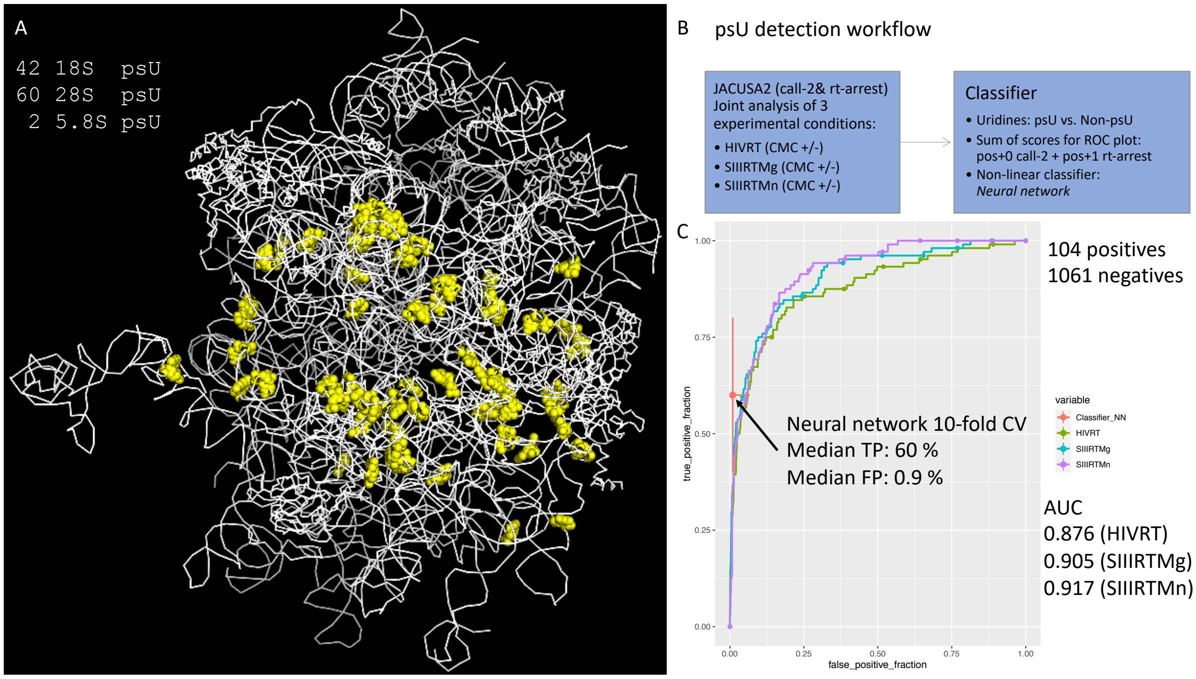


Figure 2: **Pseudouridine detection on 80S human ribosome using Illumina data from Author cite** A: Structure of the 80S ribosome (PDB file: `https://www.rcsb.org/structure/4UG0` ) with $\Psi$ positions highlighted in yellow. Annotation from [2]. B: Computational workflow to predict $\Psi$ positions at all covered U positions. One simple predictor uses the sum of scores from rt-arrest at pos+1 and call-2 at pos+0 of different RTs and a Neural network classifier uses all x scores. C: ROC plot of simple scores colored by experimental condition. Neural network performance over a 10-fold cross-validation is depicted as cross-hair and median performance values are printed out.

Our computational workflow is depicted in Figure 2B. Briefly, we compute a simple score on individual experiments by summing up the call-2 score (base substitution) at every U (position 0) and the rt-arrest score (position +1). The corresponding ROC curve is plotted in Figure 2C. Our simple neural network classifier (1 input layer with 6 nodes, 1 hidden layer with 3 nodes, 2 output nodes) reaches a mean specificity of 99%

and a mean sensitivity of 60% sensitivity on hold-out test data.

For further details, please check the corresponding Vignette in `https://github.com/dieterich-lab/JACUSA2helper`

### Accessions and BAM files

#### Zhou et al. - Pseudouridines have context-dependent mutation and stop rates in high-throughput sequencing [6]

All relevant FASTQ read files are deposited under `https://www.ncbi.nlm.nih.gov/sra/?term=SRP132322` Our respective BAM files are deposited under `https://data.dieterichlab.org`

#### Zhang et al. - Single-base mapping of m6A by an antibody-independent method [9]

All relevant FASTQ read files are deposited under `https://trace.ncbi.nlm.nih.gov/Traces/sra/?study=SRP179955`. We use the following read accession in our analysis: SRR8450805, SRR8450807, SRR8450809, SRR8450806, SRR8450808, SRR8450810

Our respective BAM files are deposited under `https://data.dieterichlab.org`

#### Meyer - DART-seq: an antibody-free method for global m6A detection [10]

All relevant FASTQ read files are deposited under `https://trace.ncbi.nlm.nih.gov/Traces/sra/?study=SRP182709`. We use the following read accession in our analysis: SRR9940470, SRR9940471, SRR9940472, SRR9940474, SRR9940475, SRR9940476

Our respective BAM files are deposited under `https://data.dieterichlab.org`

#### Pratanwanich et al. - Detection of differential RNA modifications from direct RNA sequencing of human cell lines [11]

All relevant FASTQ read files are deposited under `https://trace.ncbi.nlm.nih.gov/Traces/sra/?study=ERP124567`

### JACUSA2 parameters and downstream processing

#### Zhou et al. - Pseudouridines have context-dependent mutation and stop rates in high-throughput sequencing [6]

##### call-2 call

```
JACUSA2 Version:  2.0.0-RC22 call-2 -F 1024 -c 10 -p 10 -D -I -a D,Y,M -P1 FR-SECONDSTRAND
-P2 FR-SECONDSTRAND -r rmDup_SIIIRTMg_RC22_call2_result.out rmDup/SIIIRTMg_PLUS_CMC_global_uniq_rmdup.ba
rmDup/SIIIRTMg_MINUS_CMC_global_uniq_rmdup.bam
```

##### rt-arrest call

```
JACUSA2 Version:  2.0.0-RC22 rt-arrest -F 1024 -c 10 -p 10 -P1 FR-SECONDSTRAND -P2 FR-SECONDSTRAND
-r rmDup_SIIIRTMg_RC22_rtarrest_plain_result.out rmDup/SIIIRTMg_PLUS_CMC_global_uniq_rmdup.bam
rmDup/SIIIRTMg_MINUS_CMC_global_uniq_rmdup.bam
```

#### Zhang et al. - Single-base mapping of m6A by an antibody-independent method [9]

##### rt-arrest call

```
JACUSA2 Version:  2.0.0-RC18 rt-arrest -F 1280 -c 4 -p 10 -P1 FR-SECONDSTRAND -P2 FR-SECONDSTRAND
-r MazF_vs_cond2_FTO_RC18_rtarrest_plain_result.out SRR8450805_STARmapping_uniq_rmdup.bam,
SRR8450807_STARmapping_uniq_rmdup.bam, SRR8450809_STARmapping_uniq_rmdup.bam SRR8450806_STARmapping_uniq[illegible]
SRR8450808_STARmapping_uniq_rmdup.bam, SRR8450810_STARmapping_uniq_rmdup.bam
```

#### Meyer - DART-seq: an antibody-free method for global m6A detection [10]

##### call-2 call

```
JACUSA2 Version:  2.0.0-RC18 call-2 -F 1024 -c 4 -p 10 -D -I -a D,Y -P1 RF-FIRSTSTRAND -P2
RF-FIRSTSTRAND -r APOBEC1YTH_APOBEC1YTHmut_RC18_call2_result.out SRR9940470_STARmapping_uniq_rmdup.bam,
SRR9940471_STARmapping_uniq_rmdup.bam, SRR9940472_STARmapping_uniq_rmdup.bam SRR9940474_STARmapping_uniq[illegible]
SRR9940475_STARmapping_uniq_rmdup.bam, SRR9940476_STARmapping_uniq_rmdup.bam
```

#### Pratanwanich et al. - Detection of differential RNA modifications from direct RNA sequencing of human cell lines [11]

```
_ JACUSA2 Version:  2.0.0-RC22 call-2 -m 1 -q 1 -c 4 -p 10 -D -I -a D,Y -P1 FR-SECONDSTRAND
-P2 FR-SECONDSTRAND -r WT_vs_KO_2samp_RC22_call2_result.out HEK293T-WT-rep2.bam,HEK293T-WT-rep3.bam
HEK293T-KO-rep2.bam,HEK293T-KO-rep3.bam
   JACUSA2 Version:  2.0.0-RC22 call-2 -m 1 -q 1 -c 4 -p 10 -D -I -a D,Y -P1 FR-SECONDSTRAND
-P2 FR-SECONDSTRAND -r WT100_vs_WT0_RC22_call2_result.out HEK293T-WT-100-rep1.bam,HEK293T-WT-100-rep2.ba[illegible]
HEK293T-WT-0-rep1.bam,HEK293T-WT-0-rep2.bam
```

### Supplementary Table

#### Supplementary Table 1: m6A predictions from MazF assay (FTO+/-), DART-seq assay (YTH-domain) and Nanopore (WT/KO)

Microsoft XLSX file, 17MB, content: MazF.FTO - genomic coordinates, MazF.FTO - cDNA coordinates, DART - genomic coordinates, DART - cDNA coordinates, Nanopore - genomic coordinates, Nanopore - cDNA coordinates

#### Supplementary Table 2: $\Psi$ predictions from Zhou et al. 2018 - HIV, SIII Mg + Mn / CMC + and -

Microsoft XLSX file, 70KB, column description:
`ID` - coordinates, `SIIIRTMnCall2Score_1` - Mismatch Score position 1 (U in reference), `SIIIRTMgCall2Score_`[illegible] - Mismatch Score position 1 (U in reference), `HIVRTCall2Score_1` - Mismatch Score position 1 (U in reference), `SIIIRTMnArrestScore_2` - Arrest Score position 2 (U+1 in reference), `SIIIRTMgArrestScor`[illegible] - Arrest Score position 2 (U+1 in reference), `HIVRTArrestScore_2` - Arrest Score position 2 (U+1 in reference), `true label` - as annotated in Taoka et al. 2018, `prediction` - predicted label
